# Caspase-1-driven neutrophil pyroptosis promotes an incomplete NETosis upon *Pseudomonas aeruginosa* infection

**DOI:** 10.1101/2021.06.28.450116

**Authors:** Karin Santoni, David Pericat, Miriam Pinilla, Salimata Bagayoko, Audrey Hessel, Stephen-Adonai Leon-Icaza, Elisabeth Bellard, Serge Mazères, Emilie Doz-Deblauwe, Nathalie Winter, Christophe Paget, Jean-Philippe Girard, Céline Cougoule, Renaud Poincloux, Mohamed Lamkanfi, Emma Lefrançais, Etienne Meunier, Rémi Planès

**Author notes:** Address correspondence to (EM) and (RP), Present Address: Institute of Pharmacology and Structural Biology (IPBS), University of Toulouse, CNRS, Toulouse; France. These authors equally supervised this work (Etienne Meunier, Rémi Planès).

## Abstract

Multiple neutrophil death programs contribute to host defense against infections. Although expressing all necessary components, neutrophils specifically fail to undergo pyroptosis, a lytic form of cell death triggered by the activation of the pro-inflammatory complex inflammasome. In the light of the arm race, we hypothesized that intrinsic neutrophil pyroptosis resistance might be bypassed in response to specific microbial species. We show that *Pseudomonas aeruginosa* (*P. aeruginosa*) stimulates Caspase-1-dependent pyroptosis in human and murine neutrophils. Mechanistically, activated NLRC4 inflammasome supports Caspase-1-driven Gasdermin-D (GSDMD) activation, IL-1β cytokine release and neutrophil pyroptosis. Furthermore, GSDMD activates Peptidyl Arginine Deaminase-4 which drives an “incomplete NETosis” where neutrophil DNA fills the cell cytosol but fails crossing plasma membrane. Finally, we show that neutrophil Caspase-1 account for IL-1β production and contributes to various *P. aeruginosa* strains spread in mice. Overall, we demonstrate that neutrophils are fully competent for Caspase-1-dependent pyroptosis, which drives an unsuspected “incomplete NETosis”.

**Summary:** Neutrophils play an essential roles against infections. Although multiple neutrophil death programs contribute to host defense against infections, they fail to undergo pyroptosis, a pro-inflammatory form of cell death. Upon Infections, pyroptosis can be induced in macrophages or epithelial cells upon activation of pro-inflammatory complexes, inflammasomes that trigger Caspase-1-driven Gasdermin dependent plasma membrane lysis. In the light of host-microbe interactions, we hypothesized that yet to find microbial species might hold the capacity to overcome neutrophil resistance to inflammasome-driven pyroptosis. Among several bacterial species, we describe that the bacterium *Pseudomonas aeruginosa* specifically engages the NLRC4 inflammasome, which promotes Caspase-1-dependent Gasdermin-D activation and subsequent neutrophil pyroptosis. Furthermore, inflammasome-driven pyroptosis leads to DNA decondensation and expansion into the host cell cytosol but not to the so called Neutrophil Extracellular Trap (NET) release as DNA fails breaching the plasma membrane. Finally, *in vivo P. aeruginosa* infections highlight that Caspase-1-driven neutrophil pyroptosis is functional and is detrimental upon *P. aeruginosa* infection. Altogether, our results unexpectedly underline neutrophil competence for Caspase-1-dependent pyroptosis, a process that contributes to host susceptibility to *P. aeruginosa* infection.

## Introduction

Over the last 30 years, non-apoptotic forms of cell death have emerged as crucial processes driving inflammation, host defense against infections but also (auto) inflammatory pathologies [1].

Unique among all forms of regulated cell necrosis is the capacity of granulocyte neutrophils to undergo the process of NETosis [2]. NETosis is an antimicrobial and pro-inflammatory from of cell death that promotes the formation of extracellular web-like structures called Neutrophil Extracellular Traps (NETs) [2]. Although the importance of NETosis in host immunity to infections has been well established [2–5], NETosis dysregulation also associates to autoimmunity, host tissue damages, aberrant coagulation and thrombus that all contribute to pathology such as sepsis or autoimmune lupus [6–13].

NETosis consists in sequential steps that start with nuclear envelope disintegration, DNA decondensation, cytosolic expansion of DNA and its subsequent expulsion through plasma membrane [14]. Completion of DNA decondensation and expulsion requires various cellular effectors. Among them, neutrophil serine proteases (Neutrophil elastase, Cathepsin G, Proteinase 3) or Caspase-11 can mediate histone cleavage, which relaxes DNA tension [3,13,15–17]. In addition, granulocyte-enriched Protein arginine deaminase 4 (PAD4), citrullinates histone-bound DNA, which neutralizes arginine positive charges, thus helping nuclear DNA relaxation and decondensation [4,18,19]. Then, decondensed DNA is mixed with the neutrophil cytoplasmic granule content such as NE, CathG, PR3 and Myeloperoxidase (MPO) proteins [3,15,18]. Finally, sub-cortical actin network disassembly is required to ensure efficient DNA extrusion through the permeabilized plasma membrane [18,20].

Depending on the initial trigger, various signaling pathways such as calcium fluxes [17,18], necroptosis-associated MLKL phosphorylation [21], ROS-induced Neutrophil protease release [15] or endotoxin-activated Caspase-11 [3,5,22] all bring neutrophil into NETosis. Common to both ROS- and Caspase-11-dependent NETosis is the requirement of the pyroptosis executioner Gasdermin-D (GSDMD) cleavage by both neutrophil serine and Caspase-11 proteases, which triggers neutrophil NETosis [3,16]. Specifically, active GSDMD forms a pore on PIP2-enriched domains of the plasma and nuclear membrane of neutrophils, which ensures both IL-1-related cytokine release [23–26] and osmotic imbalance-induced DNA decondensation and expulsion [3,16].

An intriguing feature of neutrophils is that, despite GSDMD activation, they resist canonical inflammasome-induced Caspase-1-dependent cell pyroptosis upon *Salmonella* Typhimurium and *Burkholderia thaïlandensis*-activated NLRC4 inflammasome or upon Nigericin/ATP-mediated NLRP3 inflammasome activation [3,5,27,28]. However, recent studies indirectly challenged canonical pyroptosis impairment in neutrophils by showing that sterile activators of the NLRP3 inflammasome also contribute to canonical inflammasome-dependent neutrophil death and subsequent NETosis [29,30]. If it does exist specific microbial species that can promote neutrophil canonical pyroptosis has remained an open question so far.

Specifically lung infections triggered by the bacterium *Pseudomonas aeruginosa* (*P. aeruginosa*) can promote acute and chronic life-threatening infections in immunocompromised or hospitalized patients [31]. *P. aeruginosa* strains express a Type-3 Secretion System (T3SS) that allows injecting a specific set of virulence factors into host target cells, including macrophages and neutrophils [32]. T3SS-expressing *Pseudomonas aeruginosa* strains classically segregate into two mutually exclusive clades. Those expressing the bi-cistronic ADP-rybosylating and GTPase Activating Protein (GAP) virulence factor ExoS, and those expressing the lytic phospholipase of the patatin-like family, ExoU [32]. Common to most of *P. aeruginosa* strains is the expression of two other toxins, ExoY and ExoT whose functions in bacterial infection remain still unclear. All Exo toxins are injected by the T3SS into host target cells upon infections. Finally *P. aeruginosa* strains also use their T3SS to inject Flagellin into host target cells, which promotes the activation of the NAIP5-NLRC4 inflammasome and subsequent Caspase-1-driven GSDMD-dependent pyroptosis of macrophages [33–39]. Although numerous studies underlined that neutrophils are targeted by *Pseudomonas aeruginosa* virulence factors, which could promote NETosis [12,40–42], the critical effectors and their host cell targets remain extensively debated. Regarding this, defect in the Nadph oxidase (Nox2) enzyme expression has also been found to sensitive murine neutrophils to some degree of Caspase-1-driven neutrophil death upon infection with *Pseudomonas aeruginosa* [43]. This suggests that under certain conditions neutrophils might be prone to undergo Caspase-1-dependent pyroptosis. Whether such process also occurs in WT neutrophils, and its molecular as well as immune significance remain unknown.

Here, we aimed at determining if *Pseudomonas aeruginosa* but also other bacterial species could bypass neutrophil resistance to canonical pyroptosis induction as well as the key host and microbial effectors involved. We describe that the bacterial pathogen *Pseudomonas aeruginosa* uniquely triggers measurable Caspase-1-dependent pyroptosis in WT human and murine neutrophils. This requires the expression of a functional Type-3 Secretion System (T3SS) and Flagellin, but not other T3SS-derived toxins. Noticeable, deletion of Exotoxins U or S in *P. aeruginosa* entirely rewires neutrophil death towards a Caspase-1-driven pyroptosis. Specifically, *P. aeruginosa* activates the neutrophil NLRC4 inflammasome, which ensures Caspase-1-driven Gasdermin D (GSDMD) cleavage and the formation of GSDMD pores. GSDMD pores promote neutrophil pyroptosis and subsequent Peptidyl Arginine Deaminase 4 (PAD4) activation. PAD4 Citrullinates Histones, which leads to an incomplete NETosis where neutrophil DNA is decondensed and fills the host cell cytosol but is not expulsed out from the cells. Finally, using intravital microscopy on MRP8-GFP mice, we show that this incomplete NETosis occurs in neutrophils in infected lungs and that neutrophil Caspase-1 contributes both to IL-1β production and to the spread of several *P. aeruginosa* strains in mice. Overall, our results highlight that Caspase-1-dependent pyroptosis is a functional process in neutrophils which promotes a unique “incomplete NETosis”.

## Results

### Various *Pseudomonas aeruginosa* strains trigger Caspase-1-dependent and - independent neutrophil lysis

In order to determine whether neutrophils could undergo pyroptosis upon bacterial infections, we first infected WT and *Caspase-1*^-/-^ (*Casp1*^-/-^) mouse Bone Marrow Neutrophils (BMNs) with various bacterial strains, know to activate different inflammasomes in macrophages (**Fig. 1A**). We measured neutrophil lysis (LDH release) as well as IL-1β release as a hallmark of inflammasome activation. At the exception of *Staphylococcus aureus*, all bacteria triggered a Caspase-1-dependent IL-1β release, suggesting that an inflammasome-dependent response was induced in neutrophils upon various bacterial challenge (**Fig. 1A**). However, despite inducing significant neutrophil lysis, only *Pseudomonas aeruginosa* infection triggered a robust Caspase-1-dependent lysis (referred as pyroptosis) (**Fig. 1A**). Furthermore, infection of human blood neutrophils with various *Pseudomonas aeruginosa* strains (PAO1 and CHA) also triggered Caspase-1-dependent IL-1B and lysis, thus suggesting that both murine and human neutrophils are competent for Caspase-1-driven pyroptosis in response to specific bacterial species (**Fig. 1B**).

**Figure 1.**
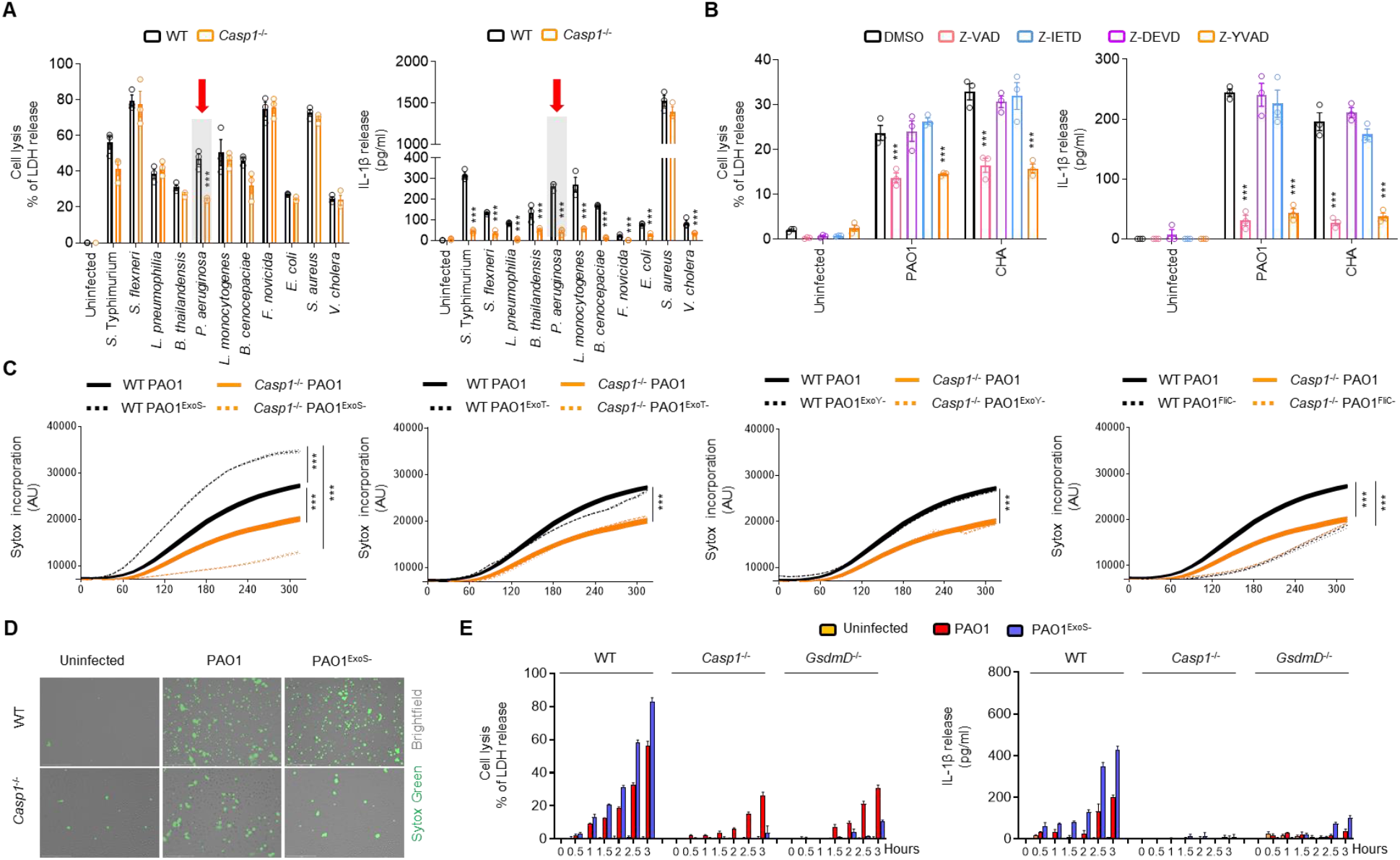
Various *Pseudomonas aeruginosa* strains trigger Caspase-1-dependent and -independent neutrophil lysis. **A.** Measure of cell lysis (release of LDH) and IL-1β release in WT or *Casp1*^-/-^ murine Bone Marrow Neutrophils (BMNs) infected for 3 hours with various bacteria at a multiplicity of infection (MOI) of 10. ***p ≤ 0.001, T-test with Bonferroni correction. Values are expressed as mean ± SEM. Graphs show one experiment representative of three independent experiments. **B.** Measure of cell lysis (release of LDH) and IL-1β release in human blood neutrophils infected for 3 hours with *Pseudomonas aeruginosa* strains PAO1 or CHA (MOI 5) in presence/absence of various Caspase inhibitors, Z-VAD (pan Caspase, 20μM), Z-YVAD (Casp1 inhibitor, 40μM), Z-DEVD (Casp3 inhibitor, 40μM), Z-IETD (Casp8 inhibitor, 40μM). ***p ≤ 0.001, T-test with Bonferroni correction. Values are expressed as mean ± SEM. Graphs show one experiment representative of three independent experiments. **C, D.** Measure of plasma membrane permeabilization and associated fluorescent microscopy images over time using SYTOX Green incorporation in WT or *Casp1*^-/-^ BMNs infected with *Pseudomonas aeruginosa* PAO1 or various isogenic mutants lacking T3SS-derived toxins ExoS, ExoY, ExoT or Flagellin (FliC) (PAO1^ExoS-^, PAO1^ExoY-^, PAO1^ExoT-^, PAO1^FliC-^). ***p ≤ 0.001, T-test with Bonferroni correction. Values are expressed as mean ± SEM. Graphs show one experiment representative of three independent experiments. Scale Bar: 150μm **E.** Measure of cell lysis (release of LDH) and IL-1β release in WT, *Casp1*^-/-^ and *GsdmD*^-/-^ murine Bone Marrow Neutrophils (BMNs) infected for the indicated times with PAO1 or its isogenic mutant PAO1^ExoS-^ at an MOI of 10. ***p ≤ 0.001, T-test with Bonferroni correction. Values are expressed as mean ± SEM. Graphs show one experiment representative of three independent experiments.

Next, we aimed at determining the means by which *P. aeruginosa* could promote Caspase-1-dependent neutrophil pyroptosis. As *P. aeruginosa* T3SS-mediated injection of toxins (ExoS, U, T, Y) and Flagellin play a major role in virulence, we hypothesized that one or many of those virulence factors might contribute to neutrophil pyroptosis. Hence, we infected murine BMNs with *P. aeruginosa* strains lacking or not expression of each toxin (PAO1^ExoS-^, PAO1^ExoT-^, PAO1^ExoY-^, PAO1^FliC-^) or deficient for the expression of the T3SS (PAO1^ExsA-^). We measured the ability of WT and *Casp1*^-/-^ neutrophils to undergo lysis (LDH release), to promote IL1β release or to exhibit plasma membrane permeabilization (SYTOX Green incorporation) upon infections with those *Pseudomonas aeruginosa* mutants.

*P. aeruginosa* strains lacking expression of T3SS or Flagellin were unable to promote a robust Caspase-1-dependent neutrophil lysis, IL-1β release and plasma membrane permeabilization, suggesting that T3SS and Flagellin are major effectors of Caspase-1-driven neutrophil death (**S1A Fig, Fig. 1C, D**).

To the contrary, strains lacking ExoY or T did not influence significantly neutrophil lysis, IL-1β release and plasma membrane permeabilization (**S1A Fig, Fig. 1C, D**). We also noticed that the infection of neutrophils with *P. aeruginosa* deficient for ExoS triggered increased lysis, membrane permabilization and IL-1β release of neutrophils (**S1A Fig, Fig. 1C, D**). In addition, PAO1^ExoS-^ -induced neutrophil lysis was completely abrogated in Caspase-1 deficient neutrophils, hence suggesting that ExoS expression might promote a Caspase-1-independent neutrophil death (**S1A Fig, Fig. 1C, D**). Caspase-1 is a central effector of macrophage lysis through the cleavage of the pyroptosis executioner Gasdermin D (GSDMD). In this context, over time measures showed that both *Casp1*^-/-^ and *GsdmD*^-/-^ neutrophils exhibited reduced LDH and IL-1β release upon *Pseudomonas aeruginosa* infection (**Fig. 1E**). Again, we remarked that PAO1^ExoS-^-induced neutrophil lysis and IL-1β release were fully CASP1- and GSDMD- dependent (**Fig. 1E**).

As 30% of *Pseudomonas aeruginosa* strains do not express ExoS but instead express the extremely lytic phospholipase toxin ExoU, we also challenged our findings by infecting neutrophils with ExoU-expressing strains (PP34^ExoU^). We observed no defect of *Casp1*^-/-^ neutrophil lysis capacity upon PP34^ExoU^ infection (**S1B, C Fig.**). However, infection of neutrophils in presence of MAFP, an inhibitor of ExoU lytic activity or with PP34^ExoU-^ or strains expressing a catalytically dead mutant of ExoU (PP34^ExoUS142A^) showed a fully Caspase-1-dependent neutrophil lysis, plasma membrane permeabilization and IL-1β release both in human and murine BMNs (**S1B, C Fig.**). This, suggests that in absence of ExoU, *P. aeruginosa* also triggers neutrophil death in a fully Caspase-1 and GSDMD-dependent manner.

Altogether, our results show the unexpected ability of Caspase-1 to promote GSDMD-dependent neutrophil lysis upon *Pseudomonas aeruginosa* infection.

### *P. aeruginosa* infection engages a canonical NLRC4-Caspase-1-Gasdermin D-dependent pyroptosis axis in neutrophils

Next, we investigated the molecular pathways by which *P. aeruginosa* promoted Caspase-1-dependent neutrophil death. The infection of WT murine neutrophils or deficient for various inflammasome sensors showed that only *Nlrc4*^-/-^ and *ASC*^-/-^ BMNs had resistance to cell lysis, IL-1β release, Caspase-1 (p20) and GSDMD (p30) processing in response to multiple strains of *P. aeruginosa* (PAO1, PAO1^ExoS-^, PP34^ExoUS142A^, CHA, CHA^ExoS-^, PP34^ExoU-^, PA14^ExoU-^) (**Fig. 2A-E, S2A, B Fig.**). Again, PAO1^ExoS-^ and PP34^ExoU-^ strains triggered exacerbated neutrophil pyroptosis, IL1β release but also Caspase-1 and GSDMD processing, a process that required NLRC4 expression (**Fig. 2A-E, S2A, B Fig.**).

**Figure 2.**
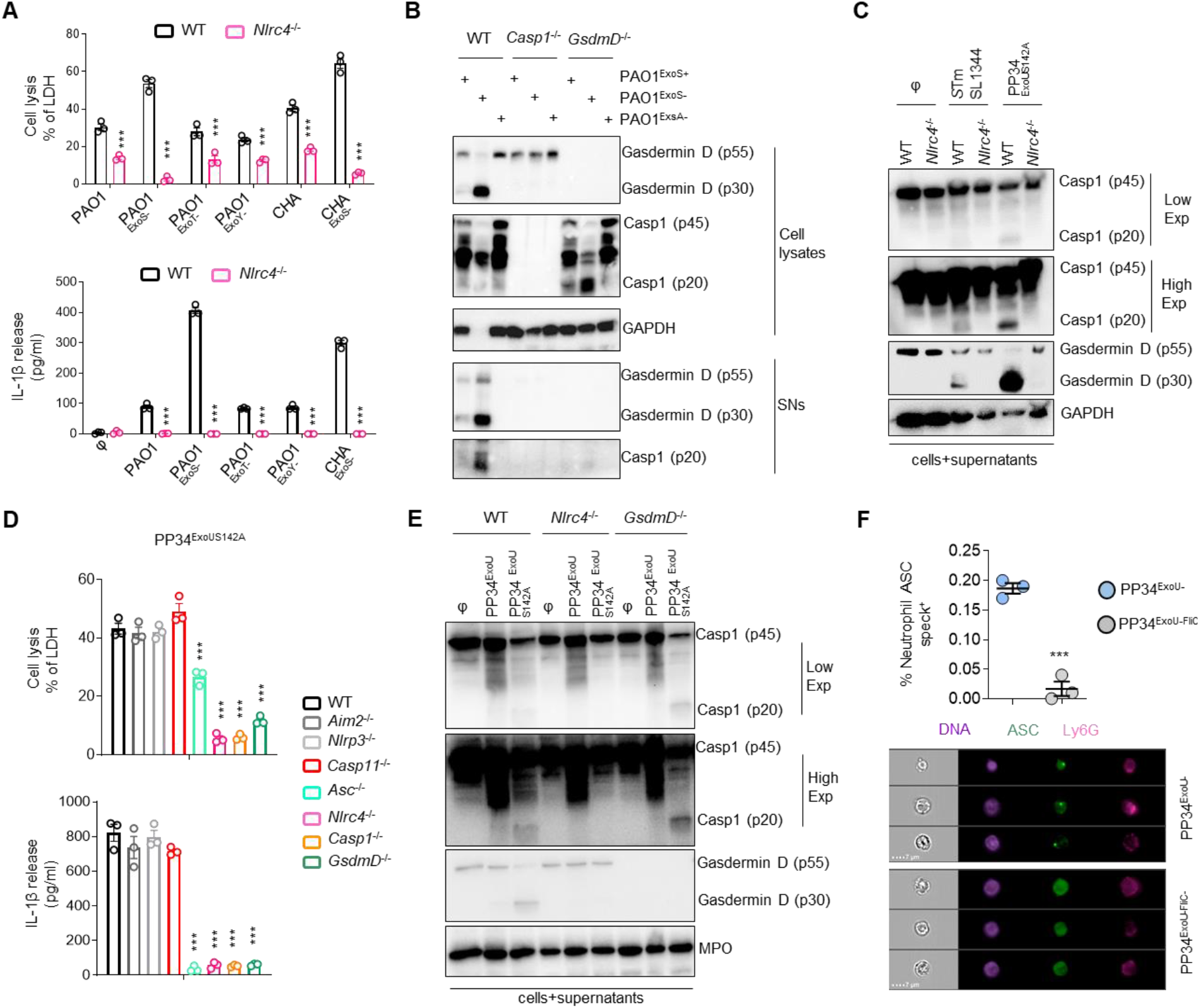
*P. aeruginosa* infection engages a canonical NLRC4-Caspase-1-Gasdermin D-dependent pyroptosis axis in neutrophils. **A.** Measure of cell lysis (release of LDH) and IL-1β release in WT or *Nlrc4*^-/-^ murine Bone Marrow Neutrophils (BMNs) infected for 3 hours with *Pseudomonas aeruginosa* PAO1 or CHA strains and their isogenic mutants lacking T3SS-derived toxins ExoS, ExoY, ExoT) at a multiplicity of infection (MOI) of 10. ***p ≤ 0.001, T-test with Bonferroni correction. Values are expressed as mean ± SEM. Graphs show one experiment representative of three independent experiments. **B.** Immunoblotting of GAPDH, pro-forms of Caspase-1 (p45) and Gasdermin-D (p55), processed Caspase-1 (p20) and Gasdermin D (p30), in WT, *Casp1*^-/-^ and *GsdmD*^-/-^ in cell lysates and cell supernatants (SNs) of BMNs infected for 3 hours with PAO1 or its isogenic mutants lacking T3SS expression (PAO1^ExsA-^) or ExoS (PAO1^ExoS-^) at a multiplicity of infection (MOI) of 10. Immunoblots show lysates and supernatants from one experiment performed three times. **C.** Immunoblotting of preforms of Caspase-1 (p45) and Gasdermin-D (p55), processed Caspase-1 (p20) and Gasdermin-D (p30) and GAPDH in WT and *Nlrc4*^-/-^ BMNs infected for 3 hours with PP34^ExoUS142A^ (MOI2) or *S*. Typhimurium (*S*.Tm, MOI 10). Immunoblots show combined lysates and supernatants from one experiment performed three times. **D.** Measure of cell lysis (release of LDH) and IL-1β release in WT or *Aim2^-/-^, Nlrp3^-/-^, Casp11^-/-^, ASC^-/-^, Nlrc4^-/-^, Casp1*^-/-^ and *GsdmD*^-/-^ murine Bone Marrow Neutrophils (BMNs) infected for 3 hours with PP34^ExoUS142A^ at a multiplicity of infection (MOI) of 2. ***p ≤ 0.001, T-test with Bonferroni correction. Values are expressed as mean ± SEM. Graphs show one experiment representative of three independent experiments. **E.** Immunoblotting of Myeloperoxidase (MPO), pro-forms of Caspase-1 (p45) and Gasdermin-D (p55), processed Caspase-1 (p20) and Gasdermin D (p30), in WT, *Nlrc4*^-/-^ and *GsdmD*^-/-^ BMNs infected for 3 hours with PP34^ExoU^ or its isogenic mutant lacking ExoU activity (PP34^ExoUS142A^) at a multiplicity of infection (MOI) of 2. Immunoblots show combined lysates and supernatants from one experiment performed three times. **F.** Imagestream experiments and quantifications of for *in vivo* formation of ASC specks in bronchoalveolar (BALs) neutrophils from ASC-Citrine mice intranasally infected with 1.10^5^ PP34^ExoU-^ or PP34^ExoU-FliC-^ for 6 hours. The gating strategy used to evaluate inflammasome activation in neutrophils was performed as follow: (i) a gate was set on cells in focus [Cells in Focus] and (ii) a sub-gate was created on single cells [Single Cells]. Then we gated first on (iii) LY6G+ Neutrophils [LY6G+] and second on (iv) ASC-citrine+ and Hoechst+ cells [Hoechst+/ASC-Citrine+] within LY6G+ population. (v) To distinguish cells with active (ASC-speck) versus inactive inflammasome (Diffuse ASC), we plotted the Intensity with the area of ASC-citrine. This strategy allows to distinguish cells with active inflammasome that were visualized and quantified. Values are expressed as mean ± SEM. Graphs show one experiment representative of two independent experiments.

Neutrophils resist NLRC4/Caspase-1-dependent pyroptosis upon *Salmonella* infection [28]. Hence, we next infected WT or *Nlrc4*^-/-^ BMNs with various bacteria (*Salmonella* Typhimurium, *Shigella flexnerii, Chromobacter violaceum, Burkholderia thailandensis*) known to trigger a NLRC4 inflammasome response and monitored for cell death and IL-1β release (**Fig. 3E**) [5,28,44–46]. None of the tested bacteria triggered a significant NLRC4-dependent neutrophil lysis of murine neutrophils although they promoted NLRC4-dependent IL-1β release and Gasdermin D processing (**S2C Fig.**).

**Figure 3.**
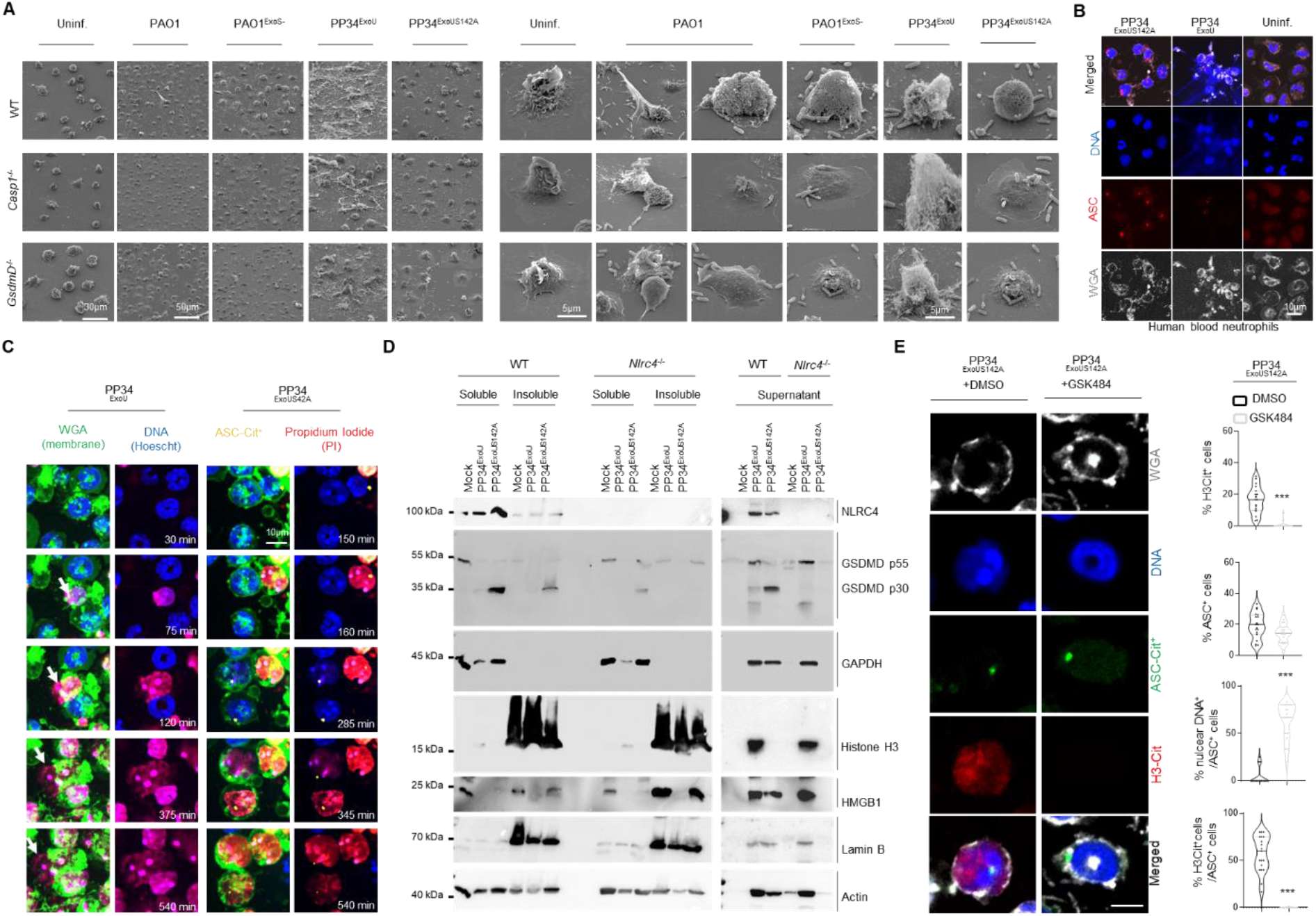
Caspase-1-induced neutrophil pyroptosis promotes PAD4-dependent incomplete NETosis. **A.** Scanning electron microscopy (SEM) observation of pyroptosis in WT, *Casp1*^-/-^ and *GsdmD*^-/-^ BMNs 3 hours after PAO1, PAO1^ExoS-^, PP34^ExoU^, PP34^ExoUS142A^ infection at an MOI of 2. Images are representative of one experiment performed 3 times. Scale bars are directly indicated in the Figure. **B.** Confocal microscopy observations of PP34^ExoU^- or PP34^ExoUS142A^-infected human blood neutrophils (MOI 2) for 3 hours harboring ASC complexes and decondensed DNA. Nucleus (blue) was stained with Hoescht; ASC is in red (anti-ASC); plasma membrane is in grey (WGA staining). Images are representative of one experiment performed 3 times. Scale bar 10μm. **C.** Representative time lapse fluorescence microscopy images of ASC-Citrine murine BMNs infected with PP34^ExoU^- or PP34^ExoUS142A^ (MOI 2) for 9 hours (540 minutes). Nucleus (blue) was stained with Hoescht; ASC is in yellow (ASC-Citrine); plasma membrane is in green (WGA staining); plasma membrane permeabilization is stained in red (cell impermanent DNA dye Propidium Iodide, PI). Images are representative of one movie out of three independent movies. Scale bar 10μm. **D.** Immunoblotting observation of Histone 3, HMGB1, Lamin B1, GAPDH, Actin, Gasdermin D (GSDMD) and NLRC4 in cellular soluble and insoluble fractions as well as in the supernatant from WT and *Nlrc4*^-/-^ murine BMNs infected with PP34^ExoU^- or PP34^ExoUS142A^ (MOI 2) for 3 hours. Immunoblots show one experiment performed three times. **E.** Confocal microscopy observations and quantifications of the percentage of cells harboring ASC complexes, H3Citrullination and nuclear/decondensed DNA in WT-ASC-Citrine^+^ BMNs infected for 3 hours with PP34^ExoUS142A^ in presence/absence of the PAD4 inhibitor GSK484 (10μM). Nucleus (blue) was stained with Hoescht; Histone-3 Citrullination is in red (Anti-H3Cit staining); plasma membrane is in grey (WGA staining). Scale bar 10μm. For quantifications, the percentage of cells with ASC complexes, nuclear DNA or positives for H3Cit (H3-Cit^+^) was determined by quantifying the ratios of cells positives for ASC speckles, nuclear DNA or H3Cit. At least 10 fields from n=3 independent experiments were analyzed. Values are expressed as mean ± SEM.

Finally, to determine if *P. aeruginosa*-induced neutrophil NLRC4 inflammasome activation also occurs *in vivo*, we infected ASC-Citrine mice with low doses (1.10^5^ CFUs) of *P*. aeruginosa strains that triggered specific NLRC4-dependent neutrophil pyroptosis, namely PP34^ExoU-^ or its isogenic mutant PP34^ExoU-/FliC-^, deficient for the expression of Flagellin (**Fig. 2D, S2D Fig.**). ImageStreamX observation of neutrophils presenting an active ASC supramolecular speck (ASC speck^+^/LY6G^+^ neutrophils) showed that PP34^ExoU-^ infection triggered inflammasome activation in neutrophils, which was reduced when mice where infected with PP34^ExoU-/FliC-^ (**Fig. 2D, S2D Fig.**). Altogether, our results show that the NLRC4/CASP1/GSDMD axis is fully functional to promote neutrophil pyroptosis in response to *P. aeruginosa* but not to various other NLRC4-activating bacteria.

### Caspase-1-induced neutrophil pyroptosis promotes PAD4-dependent incomplete NETosis

Next, we sought to determine whether Caspase-1-induced neutrophil pyroptosis could also lead to NETosis upon *P. aeruginosa* infection. We infected WT, *Casp1*^-/-^ or *GsdmD*^-/-^ murine BMNs with PAO1, PAO1^ExoS-^, PP34^ExoU^ or PP34^ExoU-^ strains and monitored for the presence of NETs using Scanning Electron Microscopy (SEM) (**Fig. 3A**). PP34^ExoU^, and to a lower extend PAO1, induced NETs in WT*, Casp1*^-/-^ and *GsdmD*^-/-^ neutrophils (**Fig. 3A**). However, we observed that neutrophil pyroptosis induced by PAO1^ExoS-^ and PP34^ExoU-^ strains failed to induce NETs (**Fig. 3A, B, S3A Fig.**). Rather, SEM and immunofluorescence experiments revealed that the fully pyroptotic strains of *P. aeruginosa* (PAO1^ExoS-^, PP34^ExoU-^, PP34^ExoUS142A^) triggered efficient DNA decondensation as well as exit from the nuclear envelop (Lamin-B1 staining) but no or few DNA release from the neutrophil plasma membrane of BMNs exhibiting an active inflammasome complex (referred as ASC specks, ASC^+^) (**Fig. 3A, B, S3A Fig.**). Further experiments using time lapse fluorescent microscopy on ASC Citrine neutrophils infected with the pyroptotic strain PP34^ExoU-^ or the NETotic strain PP34^ExoU+^ showed that both bacteria triggered efficient neutrophil DNA decondensation (**Fig. 3C, S1, 2 Movies**). Yet, pyroptotic neutrophils uniquely failed to complete DNA release out from the plasma membrane (stained with WGA) (**Fig. 3C**). NETosis drives the release of DNA and bound Histones outside from neutrophils. Hence, we reasoned that during pyroptosis, neutrophils might keep intracellularly Histone-bound DNA. Hence, immunoblotting of Histones in various neutrophil fractions (soluble, insoluble and supernatant) showed that PP34^ExoU+^-induced NETosis efficiently promoted Histone release in the extracellular medium (**Fig. 3D**). However, neither PAO1^ExoS-^ nor PP34^ExoU-^ induced Histone release, although they efficiently promoted the release of intracellular soluble and insoluble components such as GAPDH, the nuclear membrane structural component Lamin B1, the nuclear alarmin HMGB1 or NLRC4 (**Fig. 3D, S3B Fig.**). Importantly such process required NLRC4 expression (**Fig. 3D, S3B Fig.**). This suggests that Caspase-1-induced neutrophil pyroptosis can specifically promote an “incomplete NETosis” process where both DNA decondensation and release from the nuclear membrane occur, but in which DNA does not breach plasma membrane of neutrophils.

Next, we wondered about the mechanisms by which Caspase-1 promotes incomplete NETosis. Among others, Histone degradation and DNA citullination are two conserved mechanisms that promote DNA relaxation and delobulation. Neutrophil elastase and Caspase-11 promote Histone degradation and PAD4 triggers Histone citrullination. Therefore, we first explored whether Caspase-1-induced neutrophil DNA delobulation and release required PAD4-dependent citrullination. Immunoblotting and microscopy analysis of Histone citrullination showed that PP34^ExoUS142A^ and PAO1^ExoS-^ induced a robust Histone3-Citrullination in a NLRC4-, GSDMD- and CASP-1-dependent manner (**Fig. 3E, S3C-E Fig.**), a process that was also seen in human blood neutrophils (**S3C Fig.**). In order to control the specific involvement of NLRC4 at regulating Histone Citrullination in neutrophils, we used Ionomycin, a known inducer of NETosis, and observed that Ionomycin triggered NLRC4-independent Histone Citrullination in neutrophils (**S3E Fig.**). Conversely, ASC-Citrine BMNs revealed that NLRC4/CASP1/GSDMD-induced DNA decondensation required PAD4 as pharmacological inhibition of PAD4 (GSK484) abrogated both Histone citrullination as well as the DNA nuclear release (**Fig. 3E**). In addition, measure of ASC specks (ASC^+^), cell lysis (LDH release) in ASC-Citrine, WT, *Pad4*^-/-^ and *Nlrc4*^-/-^ BMNs highlighted that PAD4 was not involved in PP34^ExoUS142A^-induced NLRC4 inflammasome activation (**Fig. 3E, S3F Fig.**).

Finally, to determine if Caspase-1-induced neutrophil lysis and PAD4-dependent DNA decondensation plays a microbicidal function, we infected WT, *Pad4*^-/-^ and *Nlrc4*^-/-^ BMNs with various *Pseudomonas* strains and evaluated their cell-autonomous immune capacities. *Nlrc4*^-/-^ BMNs had improved ability to restrict PAO1, PAO1^ExoS-^ and PP34^ExoUS142A^ infection than WT and *Pad4*^-/-^ neutrophils (**S3F Fig.**), suggesting that neutrophil pyroptosis more than PAD4-driven DNA decondensation promotes neutrophil failure to restrict PAO1, PAO1^ExoS-^ and PP34^ExoUS142A^.

All in one, our results describe a pathway where NLRC4-induced neutrophil pyroptosis generates PAD4-dependent intracellular but not extracellular DNA structures.

### Neutrophil Caspase-1 contributes to IL-1β production and to *Pseudomonas aeruginosa* spread in mice

Our results showed that various *P. aeruginosa* strains trigger a Caspase-1-dependent neutrophil pyroptosis *in vitro* (**Fig. 2A**). In this context, we aimed at understanding the specific role of neutrophil Caspase-1 upon *P. aeruginosa* infection in mice. First, to determine if neutrophils can undergo Caspase-1-dependent incomplete NETosis in mice, we infected MRP8-GFP^+^ (granulocytes, including neutrophils express GFP) mice and monitored for the granulocyte death features using intravital microscopy (**Fig. 4A, S3 Movie**). Although necrotic granulocytes exhibited NETotic features (e.g. extracellular DNA) upon exposure to PP34^ExoU^, PP34^ExoUS142A^ infection led to the appearance of swelled-round necrotic granulocytes that exhibited intracellular decondensed DNA, similarly to what we observed *in vitro* (**Fig. 4A, S1, 2 Movies**). This suggests that upon lung infection, Caspase-1-induced neutrophil pyroptosis is well occurring and displays morphological and immunological distinct characteristics to NETs.

**Figure 4.**
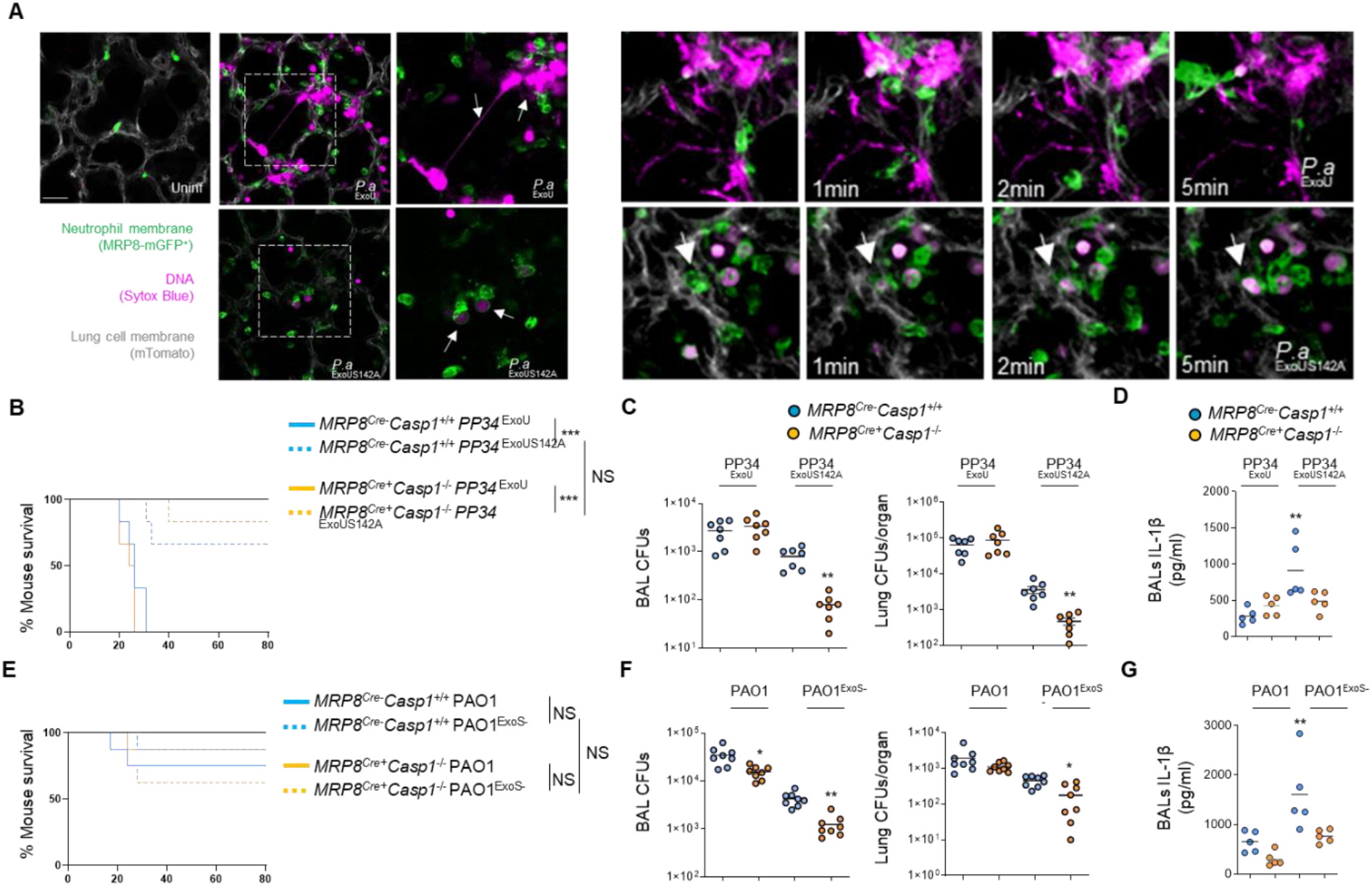
Neutrophil Caspase-1 contributes to IL-1β production and to *Pseudomonas aeruginosa* spread in mice. **A.** Intravital microscopy visualization of granulocyte death in MRP8-GFP^+^ mice infected with 2.5.10^5^ CFUs of PP34^ExoU^ (NETosis-inducing strain) or PP34^ExoUS142A^ (incomplete NETosis-inducing strain) in presence of SYTOX Blue for 10 hours. Granulocyte death was observed in infected lungs by the appearance of SYTOX blue fluorescence. Pseudo colors represent vessels (gray, mTG); Granulocytes (Green, MRP8-GFP^+^); Dead cells (Purple, SYTOX blue). Scale bar: 20μm. Data show one experiment representative of 5 independent mice. **B.** Survival of MRP8^Cre-^ *Casp1*^flox^ and MRP8^Cre+^*Casp1*^flox^ mice intranasally infected with 5.10^5^ CFUs of PP34^ExoU^ or PP34^ExoUS142A^ for 24 hours (n=6 animals per condition). Graphs represent one experiment (6 mice/group) out of three independent *in vivo* experiments. Log-rank Cox-Mantel test was used for survival comparisons. ***p ≤ 0.001. NS; Not significant. **C**. Bronchoalveolar (BAL) and lung bacterial loads (colony forming units, CFUs) in MRP8^Cre-^ *Casp1*^flox^ and MRP8^Cre+^*Casp1*^flox^ mice intranasally infected with 2.5.10^5^ CFUs of PP34^ExoU^ or PP34^ExoUS142A^ for 24 hours. Graphs represent one experiment (7 mice/group) out of three independent *in vivo* experiments; **p ≤ 0.01, Mann-Whitney analysis test. **D**. Determination of IL-1β levels in Bronchoalveolar Fluids (BALFs) MRP8^Cre-^ *Casp1*^flox^ and MRP8^Cre+^*Casp1*^flox^ mice at 10 hours after intranasal infection with 5 10^5^ CFUs (5 mice/group) of PP34^ExoU^ or PP34^ExoUS142A^. Graphs represent one experiment (5 mice/group) out of three independent *in vivo* experiments; **p ≤ 0.01, Mann-Whitney analysis test. **E.** Survival of MRP8^Cre-^ *Casp1*^flox^ and MRP8^Cre+^*Casp1*^flox^ mice intranasally infected with 1.10^7^ CFUs of PAO1^ExoS^ or PAO1^ExoS-^ for 24 hours (n=6 animals per condition). Graphs represent one experiment (6 mice/group) out of three independent *in vivo* experiments. Log-rank Cox-Mantel test was used for survival comparisons. NS; Not significant. **F**. Bronchoalveolar (BAL) and lung bacterial loads (colony forming units, CFUs) in MRP8^Cre-^ *Casp1*^flox^ and MRP8^Cre+^*Casp1*^flox^ mice intranasally infected with 5.10^6^ CFUs of PAO1^ExoS^ or PAO1^ExoS-^ for 24 hours. Graphs represent one experiment (7 mice/group) out of three independent *in vivo* experiments; *p ≤ 0.05, **p ≤ 0.01, Mann-Whitney analysis test. **G**. Determination of IL-1β levels in Bronchoalveolar Fluids (BALFs) MRP8^Cre-^ *Casp1*^flox^ and MRP8^Cre+^*Casp1*^flox^ mice at 8 hours after intranasal infection with 5. 10^6^ CFUs (5 mice/group) of PAO1^ExoS^ or PAO1^ExoS-^. Graphs represent one experiment (5 mice/group) out of three independent *in vivo* experiments; **p ≤ 0.01, Mann-Whitney analysis test.

Next, we sought to determine the importance of neutrophil Caspase-1 in response to *Pseudomonas aeruginosa* infection. Thus, we infected mice lacking CASP1 expression in the granulocytic compartment (MRP8^Cre+^*Casp1^flox^*) and their respective controls (MRP8^Cre-^*Casp1^flox^*) mice either intranasally or systemically with ExoS- or ExoU-expressing *Pseudomonas aeruginosa* (respectively PAO1^ExoS^ and PP34^ExoU^) or with their isogenic mutants PAO1^ExoS-^ and PP34^ExoU-^, both triggering a complete Caspase-1-dependent neutrophil death *in vitro*. We observed that, upon lung infections with PP34^ExoU^, MRP8^Cre+^*Casp1^flox^* mice did not show any differences in bacterial elimination, IL1β production, confirming previous work that *ExoU*-expressing *Pseudomonas* promote successful infection independently of the inflammasome pathways (**Fig. 4B-D**). To the contrary, MRP8^Cre+^*Casp1^flox^* mice infected with PAO1^ExoS^, showed a slight but significant improved bacterial elimination in Bronchoalveolar Fluids (BALF) and lungs, a phenotype that was further amplified upon infection with PAO1^ExoS-^ or PP34^ExoUS142A^ pyroptotic strains (**Fig. 4B-G**). Furthermore, IL-1β levels in BALFs were decreased in MRP8^Cre+^*Casp1^flox^* mice infected with PAO1^ExoS^, PAO1^ExoS-^ or PP34^ExoUS142A^, hence suggesting that neutrophil Caspase-1 is also a contributor of IL-1β production upon PAO1^ExoS^, PAO1^ExoS-^ or PP34^ExoUS142A^ infections (**Fig. 4D, G**).

Altogether, our results highlight that neutrophil Caspase-1 activity contributes to both IL-1β release and spreading of several *P. aeruginosa* strains.

## Discussion

Our study initially aimed at determining if neutrophil resistance to undergo Caspase-1-dependent pyroptosis can be overcome upon bacterial infection. Screening various inflammasome-activating bacteria [5,27,28], we found that *Pseudomonas aeruginosa* successfully trigger murine and human neutrophil pyroptosis through the engagement of the fully competent canonical NLRC4 inflammasome, process that requires T3SS-dependent injection of bacterial Flagellin. Although those results show a clear ability of neutrophils to undergo Caspase-1-dependent pyroptosis upon *Pseudomonas aeruginosa* infection, our investigations performed with mice specifically lacking Caspase-1 in the granulocyte compartment do suggest a minor contribution of Caspase-1 in favoring *Pseudomonas aeruginosa* spread. Related to this, previous studies showed a robust contribution of NLRC4 at promoting *Pseudomonas aeruginosa* spread in various organs, which suggests that macrophages or other NLRC4-expressing cells are stronger contributors of mouse susceptibility to *P. aeruginosa* [47,48].

Although neutrophils show intrinsic resistance to NLRC4-dependent pyroptosis (e.gs. ESCRT machinery, Caspase-1 expression levels, Ragulator pathway) [3,49–51], the unique ability of *P. aeruginosa* strains and isogenic mutants (PAO1, CHA, PAO1^ExoS-^, PP34^ExoU-^) to trigger neutrophil pyroptosis suggests that several neutrophil factors could restrict the ability of other bacteria to promote NLRC4-dependent pyroptosis. Seminal study from Zynchlynsky and colleagues found that neutrophil serine proteases could degrade the Type-3 Secretion System and flagellin virulence factors of *Shigella flexneri* [52], hence limiting their ability to hijack the neutrophil autonomous immunity and restraining *Shigella*-induced neutrophil necrosis. Similarly, upon *P. aeruginosa* infection, mouse neutrophils deficient for the NADPH oxidase enzyme Nox2 undergo increased Caspase-1-dependent pyroptosis [43]. Supporting this, Warnatsch et al., could link the extracellular Oxygen Reactive Species (ROS) localization in neutrophils exposed to *Candida albicans* to an exacerbated IL-1β production and Caspase-1 activation whereas intracellular ROS had an inhibitory effect on IL1β production and Caspase-1 activity in neutrophils [53]. Whether this could explain the capacity of the extracellularly-adapted *Pseudomonas aeruginosa* to specifically promote Caspase-1-dependent neutrophil pyroptosis but not intracellular-adapted bacterial pathogens such as *Shigella* or *Salmonella* will require further investigations.

Striking to us was the observation that expression of the key virulence factors ExoS or ExoU strongly influences neutrophils to go into NETosis whereas strains lacking ExoU or ExoS induced a complete rewiring of neutrophil death toward a Caspase-1-driven way. We hypothesize that the potent cytotoxic effect of these toxins towards various cell types, including neutrophils, may overcome inflammasome detection of *Pseudomonas aeruginosa* and triggers other neutrophil death programs [41,42,54–56]. Another non mutually exclusive guess is that these toxins directly interfere with the activation of the inflammasome pathway, thus removal of such toxins leads to an exacerbated inflammasome-response as previously reported for ExoU [39] and ExoS [57].

Although NLRC4 activation leads to Caspase-1 dependent neutrophil pyroptosis, in a pathway similar to what was previously reported in macrophage [39,48], neutrophil pyroptosis exhibits a unique feature characterized by nuclear membrane rupture, DNA decondensation and expansion within cell cytosol. One key enzyme responsible for this morphological characteristic of neutrophil pyroptosis is the Protein arginine deiminase 4 (PAD4). Indeed, similar to the process induced by various NETosis inducers, Caspase-1 also promoted PAD4-dependent Histone Citrullination, which stimulated DNA relaxation and release from the nucleus but, surprisingly, not its extracellular expulsion. Why upon Caspase-11 [3], MLKL [21], NADPH [17] or NE/CatG/Pr3 [15] stimulation but not upon Caspase-1 activation neutrophils generate two different types of DNA structures remains yet to be investigated. Recently, Thiam et al., [18] observed that pharmacological stabilization of F-actin allowed the development of this “incomplete NETosis” upon Ionomycin-exposure. Interestingly, neutrophil elastase has also been shown to degrade actin [58], hence ensuring complete NETosis process. This, suggests that efficient actin degradation and/or depolymerization may be an essential player of extracellular DNA release, which would imply that the final step of NETosis might actually be a cell-regulated process involving various controllers [14,20]. Interestingly, Chen and colleagues recently found that upon infection with *Yersinia*, murine neutrophils induce a pyroptotic program that involves virulence-inhibited innate immune sensing, hence promoting RIPK1-induced Caspase 3-dependent Gasdermin E cleavage and activation and pyroptosis [59], a process that does not trigger NETosis or “incomplete NETosis”. This further suggests that multiple Gasdermins can trigger neutrophil death through multiple molecular pathways and promote different morphological outcomes of neutrophils.

Regarding the immunological purpose of Caspase-1-induced neutrophil pyroptosis, we hypothesize that the decondensation of DNA but its conservation into the intracellular space might be a physical mean for neutrophils to trap some intracellular DAMPs, hence avoiding their passive release and a too strong exacerbation of the inflammatory response. Supporting this, we observed that DNA-bound Histones mostly remain trapped intracellularly, but not HMGB1 alarmin, both initially located in the nucleus. In the light of the recent discovery from Kayagaki and colleagues on the role for Ninjurin-1 at promoting active cell shrinkage and HMGB1/nucleosome DAMP release downstream of Gasdermin-D pores in macrophages, the use of Ninjurin-1 deficient mice are full of promises [60]. Another theoretical purpose of neutrophil “incomplete NETosis” could be the generation of “Intracellular Traps” following the “Pyroptosis-induced Intracellular Traps” model in macrophages described by Miao and colleagues. Following such speculative model, intracellular pathogens, in addition to intracellular toxic DAMPs (e.g. Histones, DNA) might remain intracellularly, hence limiting both microbial spread and pathological DAMP-dependent sterile inflammation.

All in one, our results unveil an unsuspected ability of neutrophils to undergo Caspase-1-dependent pyroptosis upon *Pseudomonas aeruginosa* infection which drives an intriguing “incomplete NETosis”, hence expanding the spectrum of neutrophil death mechanisms.

## ACKNOWLEDGEMENTS

*Nlrc4*^−/−^ mice were provided by Clare E. Bryant [61] and generated by Millenium Pharmaceutical, *GsdmD*^−/−^ mice [62] came from P. Broz (Univ of Lausanne, Switzerland), *Casp11*^−/−^ and *Casp1^−/−^/ Casp11*^−/−^ came from B. Py (ENS Lyon, France) and Junying Yuan (Harvard Med School, Boston, USA) [63,64]. Virginie Petrilli (ENS Lyon, France) provided *Nlrp3*^−/−^ mice that were generated by Fabio Martinon [65]. Thomas Henry (CIRI, Lyon, France) provided *ASC*^−/−^ and *AIM2*^−/−^ mice upon agreement with Genentech (San Francisco, Roche, USA) and. ASC-Citrine (#030744) and *Pad4*^−/−^ (#030315) mice came from Jaxson Laboratory (USA) and were generated by Douglas T Golenbock (University of Massachusetts Medical School, USA) and Kerri Mowen (The Scripps Research Institute, USA) respectively. MRP8^*Cre*^/*Casp1*^flox^ mice are provided by Natalie Winter (INRAE Tours Nouzilly, France) and were generated by crossing MRP8^*Cre*^ (Jackson # 021614) mice with *Caspase1^flox^* mice generated by Mohamed Lamkanfi (Univ. of Ghent, Belgium)[66]. MRP8^CreGFP^ and mTmG mice were obtained from Jackson laboratories and generated respectively by Emmanuelle Passegue (UCSF, USA) and Liqun Luo, (Stanford University, USA). *Pseudomonas aeruginosa* strains were a kind gift of Ina Attrée (CNRS, Grenoble, France) and Julien Buyck (Univ. of Poitiers, France). Authors also acknowledge the animal facility and Cytometry/microscopy platforms of the INFINITY, CBI and IPBS institutes and particularly Valerie Duplan-Eche for Imagestream acquisition and analysis. This project was funded by grants from the Fonds de Recherche en Santé Respiratoire - Fondation du Souffle to EL, ATIP-Avenir program, FRM “Amorçage Jeunes Equipes” (AJE20151034460) and the ERC (StG INFLAME 804249) to EM, the European Society of Clinical Microbiology and Infectious Diseases (ESCMID, 2020) to RP, Invivogen-CIFRE collaborative PhD (to MP) and post-doctoral fellowships (to RP) and from Mali and Campus France cooperative agencies to SB. Funders had not interest and regard in the conduct of the project.

## AUTHOR CONTRIBUTIONS

RP and EM designed the experiments and acquired funding. RP, KS and EM wrote the manuscript. RP and KS performed the experiments with the help of DP, MP, SB, PJB, AH, SALI, CC. Specifically RP and RP performed SEM experiments, SM, EB and EL set up and performed intravital mouse experiments, JPG, ML and NW provided essential reagents, tools and inputs for the conduct of the project. EM, RP and KS supervised the entire study.

## Supplemental Figures

**S1 Movie.** Intravital microscopy visualization of granulocyte death in MRP8-GFP^+^ mice infected with 2.5.10^5^ CFUs of PP34^ExoU^ (NETosis-inducing strains) or PP34^ExoUS142A^ (incomplete NETosis-inducing strain) in presence of SYTOX Blue for 10 hours. Granulocyte death was observed in infected lungs by the appearance of SYTOX blue fluorescence. Pseudo colors represent vessels (gray, mTG); Granulocytes (Green, MRP8-GFP+); Dead cells (Purple, SYTOX blue). Scale bar: 20μm.

**S2 Movie.** Time Lapse Fluorescence microscopy of ASC-Citrine murine BMNs infected with PP34^ExoU^ (MOI 2) for 9 hours (540 minutes). Nucleus (blue) was stained with Hoescht; ASC is in yellow (ASC-Citrine); plasma membrane is in green (WGA staining); plasma membrane permeabilization is stained in red (cell impermanent DNA dye Propidium Iodide, PI).

**S3 Movie.** Time Lapse Fluorescence microscopy of ASC-Citrine murine BMNs infected with PP34^ExoUS142^ (MOI 2) for 9 hours (540 minutes). Nucleus (blue) was stained with Hoescht; ASC is in yellow (ASC-Citrine); plasma membrane is in green (WGA staining); plasma membrane permeabilization is stained in red (cell impermanent DNA dye Propidium Iodide, PI).

**Supplemental Figure 1.**
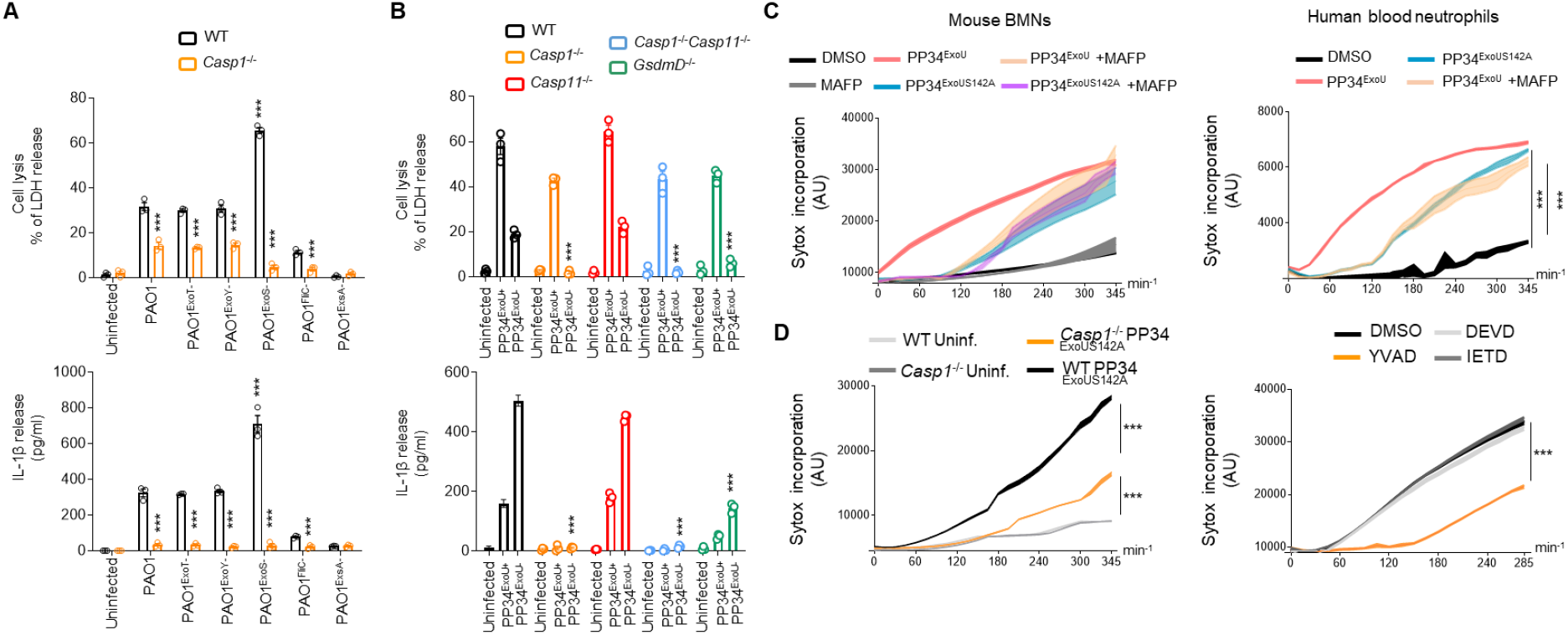
Multiple *P. aeruginosa* strains triggers Caspase-1-dependent pyroptosis in neutrophils. **A.** Measure of cell lysis (release of LDH) and IL-1β release in WT or *Casp1*^-/-^ murine Bone Marrow Neutrophils (BMNs) infected for 3 hours with *Pseudomonas aeruginosa* PAO1 and its isogenic mutants lacking T3SS-derived toxins ExoS, ExoY, ExoT or Flagellin (FliC) (PAO1^ExoS-^, PAO1^ExoY-^, PAO1^ExoT-^, PAO1^FliC-^) at a multiplicity of infection (MOI) of 10. ***p ≤ 0.001, T-test with Bonferroni correction. Values are expressed as mean ± SEM. Graphs show one experiment representative of three independent experiments. **B.** Measure of cell lysis (release of LDH) and IL-1β release in WT, *Casp1^-/-^, Casp1^-/-^ Casp11^-/-^, Casp11*^-/-^ and *GsdmD*^-/-^ murine Bone Marrow Neutrophils (BMNs) infected for 3 hours with *Pseudomonas aeruginosa* PP34^ExoU^ and its isogenic mutant PP34^ExoU-^ at a multiplicity of infection (MOI) of 2. ***p ≤ 0.001, T-test with Bonferroni correction. Values are expressed as mean ± SEM. Graphs show one experiment representative of three independent experiments. **C.** Measure of plasma membrane permeabilization over time using SYTOX Green incorporation in murine BMNs or in human blood neutrophils infected with PP34^ExoU^ and its isogenic catalytically inactive mutant PP34^ExoUS142A^ at a multiplicity of infection (MOI) of 2 in presence/absence of the phospholipase inhibitor MAFP (20μM). ***p ≤ 0.001, T-test with Bonferroni correction. Values are expressed as mean ± SEM. Graphs show one experiment representative of three independent experiments. **D.** Measure of plasma membrane permeabilization over time using SYTOX Green incorporation in murine WT and *Casp1*^-/-^ BMNs or in human blood neutrophils infected with PP34^ExoU^ and its isogenic catalytically inactive mutant PP34^ExoUS142A^ at a multiplicity of infection (MOI) of 2 in presence/absence of various Caspase inhibitors, Z-YVAD (Casp1 inhibitor, 20μM), Z-DEVD (Casp3 inhibitor, 20μM), Z-IETD (Casp8 inhibitor, 20μM). ***p ≤ 0.001, T-test with Bonferroni correction. Values are expressed as mean ± SEM. Graphs show one experiment representative of three independent experiments.

**Supplemental Figure 2.**
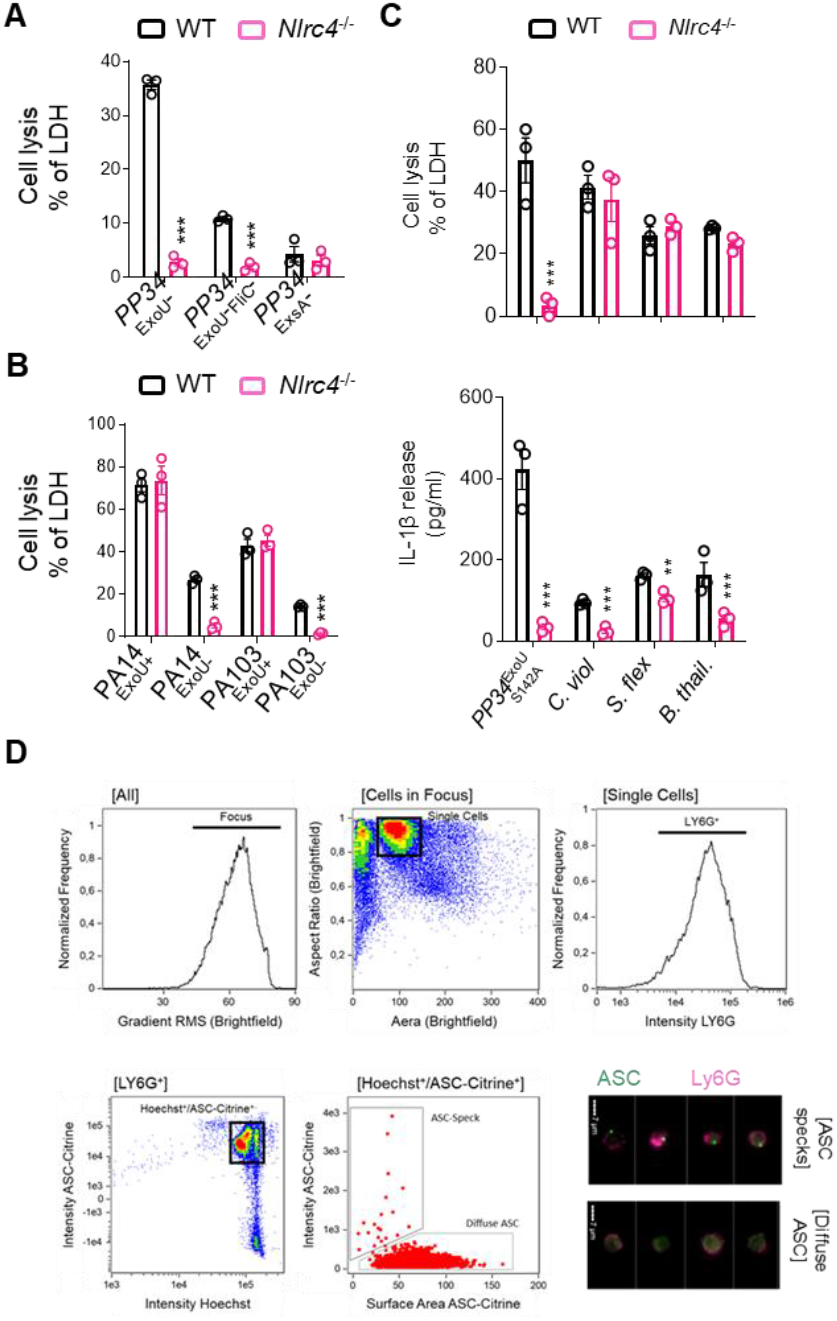
*P. aeruginosa* uniquely triggers a NLRC4-dependent pyroptosis in neutrophils. **A, B.** Measure of cell lysis (release of LDH) in WT or *Nlrc4*^-/-^ murine Bone Marrow Neutrophils (BMNs) infected for 3 hours with various ExoU-expressing *Pseudomonas aeruginosa* strains (PP34, PA14, PA103) or their isogenic mutants lacking T3SS-derived ExoU, Flagellin (FliC) or T3SS expression (ExsA) at a multiplicity of infection (MOI) of 2 (PP34) or 10 (PA14, PA103). ***p ≤ 0.001, T-test with Bonferroni correction. Values are expressed as mean ± SEM. Graphs show one experiment representative of three independent experiments. **C.** Measure of cell lysis (release of LDH) and IL-1β release in WT and *Nlrc4*^-/-^ BMNs infected for 3 hours with PP34^ExoUS142A^ (MOI 2), *Salmonella* Typhimurium (*S*.Tm SL1344, MOI 10), *Chromobacter violaceum* (*C. violaceum*, MOI 20), *Shigella flexneri* (*S. flexneri*, MOI 40) or *Burkholderia thailandensis* (*B. thailandensis*, MOI 40). ***p ≤ 0.001, T-test with Bonferroni correction. Values are expressed as mean ± SEM. Graphs show one experiment representative of three independent experiments. **D.** Gating strategy used to evaluate inflammasome activation in neutrophils was performed as follow: (i) a gate was set on cells in focus [Cells in Focus] and (ii) a sub-gate was created on single cells [Single Cells]. Then we gated first on (iii) LY6G^+^ Neutrophils [LY6G^+^] and second on (iv) ASC-citrine^+^ and Hoechst^+^ cells [Hoechst^+^/ASC-Citrine^+^] within LY6G^+^ population. (v) To distinguish cells with active (ASC-speck) versus inactive inflammasome (Diffuse ASC), we plotted the Intensity with the area of ASC-citrine. This strategy allow to distinguish cells with active inflammasome that were visualized and quantified. Values are expressed as mean ± SEM. Graphs show one experiment representative of two independent experiments.

**Supplemental Figure 3.**
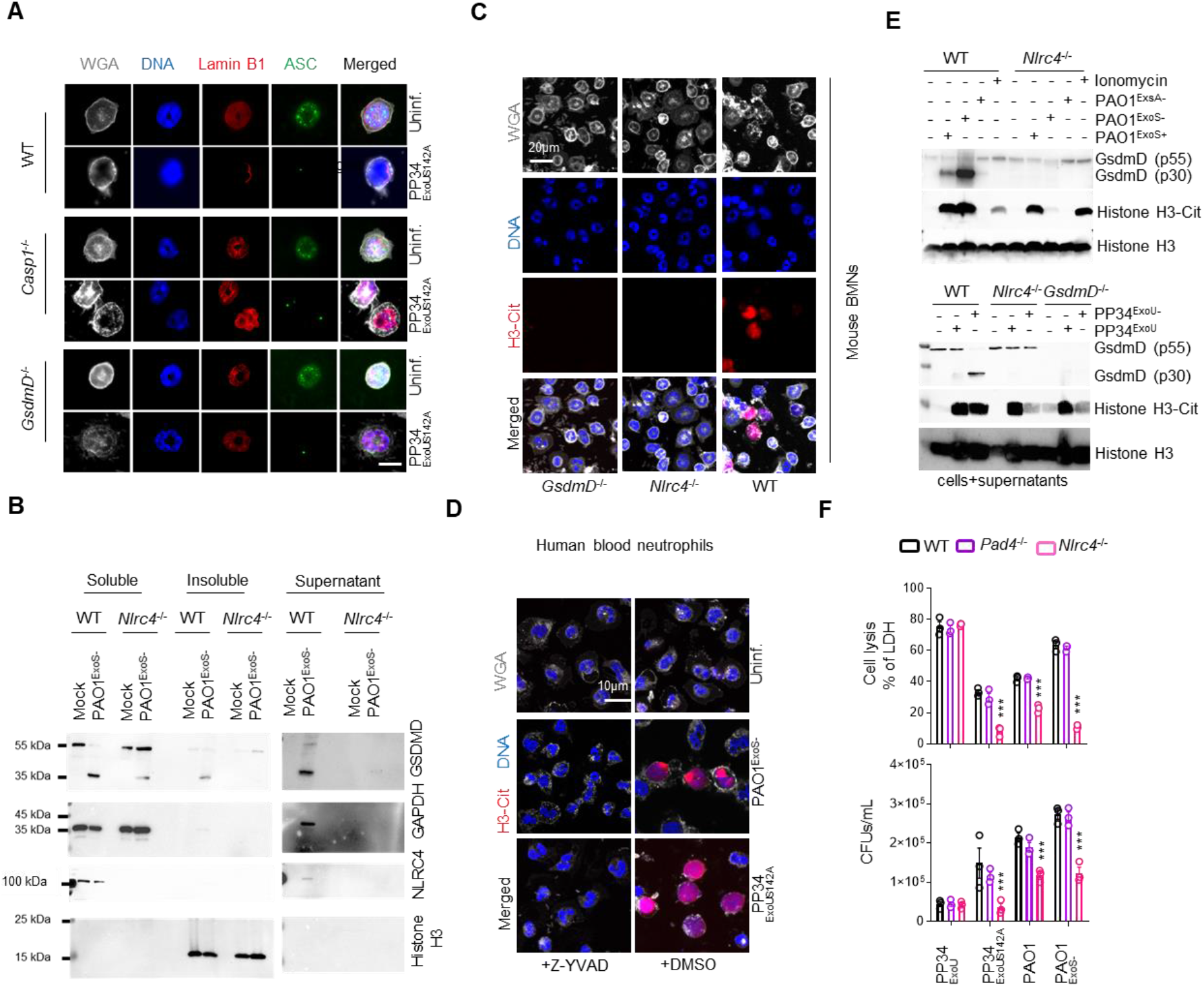
NLRC4-dependent neutrophil pyroptosis promotes an incomplete NETosis. **A.** Confocal microscopy observations of WT, *Casp1*^-/-^ and *GsdmD*^-/-^ BMNs infected for 3 hours with PP34^ExoUS142A^ and harboring ASC complexes, decondensed DNA and nuclear membrane (LaminB1). Nucleus (blue) was stained with Hoescht; LaminB1 is in red (anti LaminB1); ASC is in Green (anti-ASC); plasma membrane is in grey (WGA staining). Scale bar 10μm. Images are representative of one experiment performed three times with at least 200 neutrophils observed/ experiment. **B.** Immunoblotting observation of Histone 3, HMGB1, Lamin B1, GAPDH, Actin, Gasdermin D (GSDMD) and NLRC4 in cellular soluble and insoluble fractions as well as in the supernatant from WT and *Nlrc4*^-/-^ murine BMNs infected with PAO1^ExoS-^ (MOI 10) for 3 hours. Immunoblots show one experiment performed two times. **C.** Confocal microscopy observations of WT, *Nlrc4*^-/-^ and *GsdmD*^-/-^ BMNs infected for 3 hours with PP34^ExoUS142A^ and harboring Citrullinated Histone 3 (H3-Cit), decondensed DNA and nuclear membrane (LaminB1). Nucleus (blue) was stained with Hoescht; Citrullinated Histone-3 is in red (anti H3-Cit); plasma membrane is in grey (WGA staining). Scale bar 10μm. Images are representative of one experiment performed three times with at least 150 neutrophils observed/ experiment. **D.** Confocal microscopy observations of human blood neutrophils infected for 3 hours with PAO1^ExoS-^ (MOI 10) or PP34^ExoUS142A^ (MOI 2) in presence/absence of Caspase-1 inhibitor Z-YVAD (20μM) and harboring Citrullinated Histone 3 (H3-Cit), decondensed DNA and nuclear membrane (LaminB1). Nucleus (blue) was stained with Hoescht; Citrullinated Histone-3 is in red (anti H3-Cit); plasma membrane is in grey (WGA staining). Scale bar 10μm. Images are representative of one experiment performed three times with at least 150 neutrophils observed/ experiment. **E.** Immunoblotting of Citrullinated Histone 3 (H3Cit), total Histone 3 and preformed and cleaved Gasdermin-D (p55/p30) in WT, *Nlrc4*^-/-^ and *GsdmD*^-/-^ BMNs treated with Ionomycin (10μM, 3 hours) or infected for 3 hours with PAO1, PAO1^ExoS-^, PAO1^ExsA-^ (MOI 10) or with PP34^ExoU^, PP34^ExoU-^ (MOI 2). Immunoblots show combined lysates and supernatants from one experiment performed three times. **F.** Measure of cell lysis (release of LDH) and bacterial elimination (Colony-forming Units, CFUs) in WT, *Pad4*^-/-^ and *Nlrc4*^-/-^ BMNs infected for 3 hours with PP34^ExoU^ (MOI2), PP34^ExoUS142A^ (MOI2), PAO1 (MOI 10) or PAO1^ExoS-^ (MOI 10). ***p ≤ 0.001, T-test with Bonferroni correction. NS: Not significant. Values are expressed as mean ± SEM. Graphs show one experiment representative of three independent experiments.

## Methods

All reagents, concentrations of use and their references are listed in **Table S1**

### Mice

*Nlrc4*^−/−^ [61], *Nlrp3*^−/−^ [65], *ASC^−/−^, Casp11*^−/−^ [63,64], *Casp1*^−/−^*Casp11*^−/−^ [63,64], *GsdmD^−/−^, Aim2^−/−^, Pad4*^−/−^,*MRP8*^Cre+^*GFP*^+^, *MRP8*^Cre+^*Casp1*^flox^ were generated and described in previous studies. Mice were bred at the IPBS (Toulouse, France) and INRAE (Tours Nouzilly, France) animal facilities in agreement to the EU and French directives on animal welfare (Directive 2010/63/EU). Charles Rivers provided WT C57BL/6 mice. Mice experiments are under legal authorizations APAFIS#8521-2017041008135771 and APAFIS#12812-2018031218075551, according to the local, French and European ethic laws.

### MRP8^Cre^*Casp1*^flox^ mice genotyping

*Casp1^flox/flox^* mice were kindly provided by Mo Lamkanfi and were crossed to MRP8^Cre^ mice to generate MRP8^Cre^*Casp1*^flox^. *Caspase-1* genotyping was performed using Primer Fw: CGAGGGTTGGAGCTCAAGTTGACC and Primer Rv: CACTTTGACTTCTCTAAGGACAG. *Cre* genotyping was performed using Primers Fw: CGCCGTAAATCAATCGATGAGTTGCTTC and Primers Rv: GATGCCGGTGAACGTGCAAAACAGGCTC.

### Bacterial cultures

*P. aeruginosa* strains (PAO1, CHA, PP34, PA14, PA103) and their isogenic mutants were grown overnight in Luria Broth (LB) medium at 37°C with constant agitation. Bacteria were sub-cultured the next day by diluting overnight culture 1:25 and grew until reaching an optical density (OD) O.D.600 of 0.6 – 0.8. Bacterial strains and their mutants are listed in **Table S1**.

### Bacterial KO generation and complementation

The knockout vector pEXG2 was constructed and used based on the protocol described by Rietsch et al. [67] with the following modifications. Briefly, 700-bp sequences of the flanking regions of the selected gene were amplified by PCR with Q5 high fidelity polymerase (New England Biolabs). Then, the flanking regions were gel purified and inserted into pEXG2 plasmid by Gibson assembly [68]. The assembled plasmid was directly transformed into competent SM10λpir using Mix&Go competent cells (Zymo Research Corporation) and plated on selective LB plates containing 50 μg/mL kanamycin and 15 μg/mL gentamicin. The resulting clones were sequenced, and mating was allowed for 4 h with PAO1 strain at 37°C. The mated strains were selected for single cross over on plates containing 15 μg/mL gentamicin and 20 μg/mL Irgasan (removal of E.coli SM10 strains). The next day, some clones were grown in LB for 4 hours and streaked on 5% sucrose LB plates overnight at 30°C. P. aeruginosa clones were then checked by PCR for mutations. All primers were designed with Snapgene software (GSL Biotech LLC).

### Mice infections

Age and sex-matched animals (5–8 weeks old) per group were infected intranasally with 5.10^5^ (lethal doses) or 2.5.10^5^ CFUs of PP34^ExoU^/PP34^ExoUS142A^ or with 1.10^7^ CFUs (lethal doses) or 5.10^6^ CFUs of PAO1/PAO1^ExoS-^ strains suspended in 25μL of PBS. Animals were sacrificed at indicated times after infection and bronchoalveolar fluids (BALFs) and lungs were recovered. When specified, bacterial loads (CFU plating), cytokine levels (ELISA) were evaluated. No randomization or blinding were done.

### Intravital microscopy experiments

We relied on the previously published lung intravital microscopy method using an intercoastal thoracic window [69,70], adapted at the IPBS CNRS-University of Toulouse TRI platform.

MRP8-mTmG mice (8-12 weeks old) were infected intratracheally with 5.10^5^ CFUs of *P. aeruginosa* ExoU or ExoU^S142A^ strains resuspended in 50μL of PBS and imaged 6 to 8 hours after infection. 50μL of 50μM solution of SYTOX blue (Life Technologies) was injected both intravenously (retroorbital) and intratracheally just before imaging to visualize extracellular DNA.

Next, mice were anesthetized with ketamine and xylazine and secured to a microscope stage. A small tracheal cannula was inserted, sutured and attached to a MiniVent mouse ventilator (Harvard Apparatus). Mice were ventilated with a tidal volume of 10 μl of compressed air (21% O_2_) per gram of mouse weight, a respiratory rate of 130-140 breaths per minute, and a positive-end expiratory pressure of 2-3 cm H_2_O. Isoflurane was continuously delivered to maintain anesthesia and 300 μl of 0.9% saline solution were i.p. administered in mice every hour for hydration. Mice were placed in the right lateral decubitus position and a small surgical incision was made to expose the rib cage. A second incision was then made into the intercostal space between ribs 4 and 5, through the parietal pleura, to expose the surface of the left lung lobe. A flanged thoracic window with an 8 mm coverslip was inserted between the ribs and secured to the stage using a set of optical posts and a 90° angle post clamp (Thor Labs). Suction was applied to gently immobilize the lung (Dexter Medical). Mice were placed in 30°C heated box during microscopy acquisition to maintain the body temperature and the 2-photon microscope objective was lowered over the thoracic window. Intravital imaging was performed using a Zeiss 7MP upright multi-photon microscope equipped with a 20×/1.0 objective and a Ti-Sapphire femtosecond laser, Chameleon-Ultra II (Coherent Inc.) tuned to 920 nm. SYTOX Blue, GFP and Tomato emission signals were detected thanks to the respective bandpass filters: Blue (SP485), Green (500-550) and Red (565-610). Images were analyzed using Imaris software (Bitplane) and Zen (Zeiss).

### Isolation of primary murine neutrophils

Murine Bone marrow cells were isolated from tibias and femurs, and neutrophils were purified by positive selection using Anti-Ly-6G MicroBead Kit (Miltenyi Biotech) according to manufacturer’s instructions. This process routinely yielded >95% of neutrophil population as assessed by flow cytometry of Ly6G^+^/CD11b^+^ cells.

### Isolation of primary human neutrophils

Whole blood was collected from healthy donors by the “Ecole française du sang” (EFS, Tolouse Purpan, France) in accordance with relevant guidelines. Written, informed consent was obtained from each donor. Neutrophils were then isolated by negative selection using MACSxpress® Whole Blood Human Neutrophil Isolation Kit (Miltenyi Biotech) according to manufacturer’s instructions. Following isolations cells were centrifuged 10 min at 300 g and red blood cells were eliminated using Red blood cells (RBC) Lysis Buffer (BioLegend). This procedure gives >95% of neutrophil population as assessed by flow cytometry of CD15+/CD16+ cells. License to use human samples is under legal agreement with the EFS; contract n° 21PLER2017-0035AV02, according to Decret N° 2007-1220 (articles L1243-4, R1243-61).

### Cell plating and treatment of Neutrophils

Following isolation, Neutrophils were centrifugated for 10 min at 300 g and pellet was resuspendent in serum free OPTI-MEM medium. Absolute cell number was determined with automated cell counter Olympus R1 with trypan blue cell death exclusion method (typically living cells represent >70% of cell solution) and cell density was adjusted at 10^6^ / mL by adding OPTI-MEM culture medium. Neutrophils were then plated in either 96 well plates, 24 well plates or 6 well plates with 100 μL (10^5^ cells), 500 μL (5.10^5^ cells) or 2 mL (2.10^6^ cells) respectively. When indicated cells were incubated with chemical inhibitors Z-VAD-fmk (20 μM), Y-VAD-fmk (40 μM), Z-DEVD-fmk (40μM), Z-IETD-fmk (40μM), GSK484 (10 μM), as indicated in each experimental setting. Neutrophils were infected with various bacterial strains and multiplicity of infections (M.O.I.) as indicated.

### Kinetic analysis of Neutrophil’s permeability by SYTOX Green incorporation assay

Cells are plated at density of 1 x 10^5^ per well in Black/Clear 96-well Plates in OPTI-MEM culture medium supplemented with SYTOX-Green dye (100ng/mL) and infected/treated as mentioned in figure legend. Green fluorescence are measured in real-time using Clariostar plate reader equipped with a 37°C cell incubator.

### ELISA and plasma membrane lysis tests

Cell death was measured by quantification of the lactate dehydrogenase (LDH) release into the cell supernatant using LDH Cytotoxicity Detection Kit (Takara). Briefly, 100 μL cell supernatant were incubated with 100 μL LDH substrate and incubated for 15 min. The enzymatic reaction was stopped by adding 50 μL of stop solution. Maximal cell death was determined with whole cell lysates from unstimulated cells incubated with 1% Triton X-100. Human and mouse IL-1β secretion was quantified by ELISA kits (Thermo Fisher Scientific) according to the manufacturer’s instructions.

### Preparation of neutrophil lysates and supernatant for immunoblot

At the end of the treatment 5 mM of diisopropylfluorophosphate (DFP) cell permeable serine protease inhibitor was added to cell culture medium. Cell’ Supernatant was collected and clarified from non-adherent cells by centrifugation for 10 min at 300 g. Cell pellet and adherent cells were lysed in 100 μL of RIPA buffer (150 mM NaCl, 50 mM Tris-HCl, 1% Triton X-100, 0.5% Na-deoxycholate) supplemented with 5 mM diisopropylfluorophosphate (DFP) in addition to the protease inhibitor cocktail (Roche). Cell scrapper was used to ensure optimal recovery of cell lysate. Collected cell lysate was homogenized by pipetting up and down ten times and supplemented with laemli buffer (1X final) before boiling sample for 10 min at 95°C. Soluble proteins from cell supernatant fraction were precipitated as described previously [71]. Precipitated pellet was then resuspended in 100 μL of RIPA buffer plus laemli supplemented with 5 mM diisopropylfluorophosphate (DFP) and protease inhibitor cocktail (Roche) and heat denaturated for 10 min at 95°C. Cell lysate and cell supernatant fraction were then analysed by immunoblot either individually or in pooled sample of lysate plus supernatant (equal vol/vol).

### Treatment of Neutrophils for Immunofluorescences

5.10^5^ Cells were plated on 1.5 glass coverslips in 24 well plate and infected/treated as described above. At the end of the assay, cell supernatant was removed and cells were fixed with a 4% PFA solution for 10 min at 37°C. PFA was then removed and cells were washed 3 times with HBSS. When desired, plasma membrane was stained with Wheat Germ Agglutinin, Alexa Fluor™ 633 Conjugate (ThermoFisher Scientifique) at 1/100^th^ dilution in HBSS, and incubated for 30 min under 100 rpm orbital shaking conditions. Then cells were washed with HBSS and processed for further staining. Permeabilization was performed by incubating cells for 10 min in PBS containing 0.1% Triton X-100. To block unspecific binding of the antibodies cells are incubated in PBS-T (PBS+ 0.1% Tween 20), containing 2% BSA, 22.52 mg/mL glycine in for 30 min. 3 washes with PBS-T was performed between each steps. Primary antibodies staining was performed overnight at 4°C in BSA 2% - Tween 0.1% - PBS (PBS-T) solution. Coverslips were washed three times with PBS-T and incubated with the appropriate fluor-coupled secondary antibodies for 1 hour at room temperature. DNA was counterstained with Hoechst. Cells were then washed three times with PBS and mounted on glass slides using Vectashield (Vectalabs). Coverslips were imaged using confocal Zeiss LSM 710 (INFINITY, Toulouse) or Olympus Spinning disk (Image core Facility, IPBS, Toulouse) using a 40x or a 63x oil objective. Unless specified, for each experiment, 5-10 fields (~50-250 cells) were manually counted using Image J software.

### Scanning Electron Microscopy experiments

For scanning electron microscopy observations, cells were fixed with 2.5% glutaraldehyde in 0.2M cacodylate buffer (pH 7.4). Preparations were then washed three times for 5min in 0.2M cacodylate buffer (pH 7.4) and washed with distilled water. Samples were dehydrated through a graded series (25 to 100%) of ethanol, transferred in acetone and subjected to critical point drying with CO2 in a Leica EM CPD300. Dried specimens were sputter-coated with 3 nm platinum with a Leica EM MED020 evaporator and were examined and photographed with a FEI Quanta FEG250.

### ImageStreamX

Cells isolated from peritoneal washes were pelleted by centrifugation (10 min at 300 g). Neutrophils were stained prior to fixation with anti-Ly6G (APC-Vio770, Miltenyi-Biotec Clone: REA526 | Dilution: 1:50) in MACS buffer (PBS-BSA 0,5%-EDTA 2mM) in presence of FC block (1/100) and Hoechst (1 μM). Then, cells were fixed in 4% PFA. Data were acquired on ImageStreamXMKII (Amnis) device (CPTP Imaging and Cytometry core facility) and analyzed using IDEAS software v2.6 (Amnis). The gating strategy used to evaluate inflammasome activation in neutrophils was performed as follow: (i) a gate was set on cells in focus [Cells in Focus] and (ii) a sub-gate was created on single cells [Single Cells]. Then we gated first on (iii) LY6G+ Neutrophils [LY6G+] and second on (iv) ASC-citrine+ and Hoechst+ cells [Hoechst+/ASC-Citrine+] within LY6G+ population. (v) To distinguish cells with active (ASC-speck) versus inactive inflammasome (Diffuse ASC), we plotted the Intensity with the area of ASC-citrine. This strategy allow to distinguish cells with active inflammasome that were visualized and quantified (**S2 Fig.**).

### Statistical tests used

Statistical analysis was performed with Prism 8.0a (GraphPad Software, Inc.). Otherwise specified, data are reported as mean with SEM. T-test with Bonferroni correction was chosen for comparison of two groups. For *in vivo* mice experiments and comparisons we used Mann-Whitney tests and mouse survival analysis were performed using log-rank Cox-Mantel test. P values are shown in figures with the following meaning; NS non-significant and Significance is specified as *p ≤ 0.05; **p ≤ 0.01, ***p ≤ 0.001.

---

**Supplemental 1 Table.**
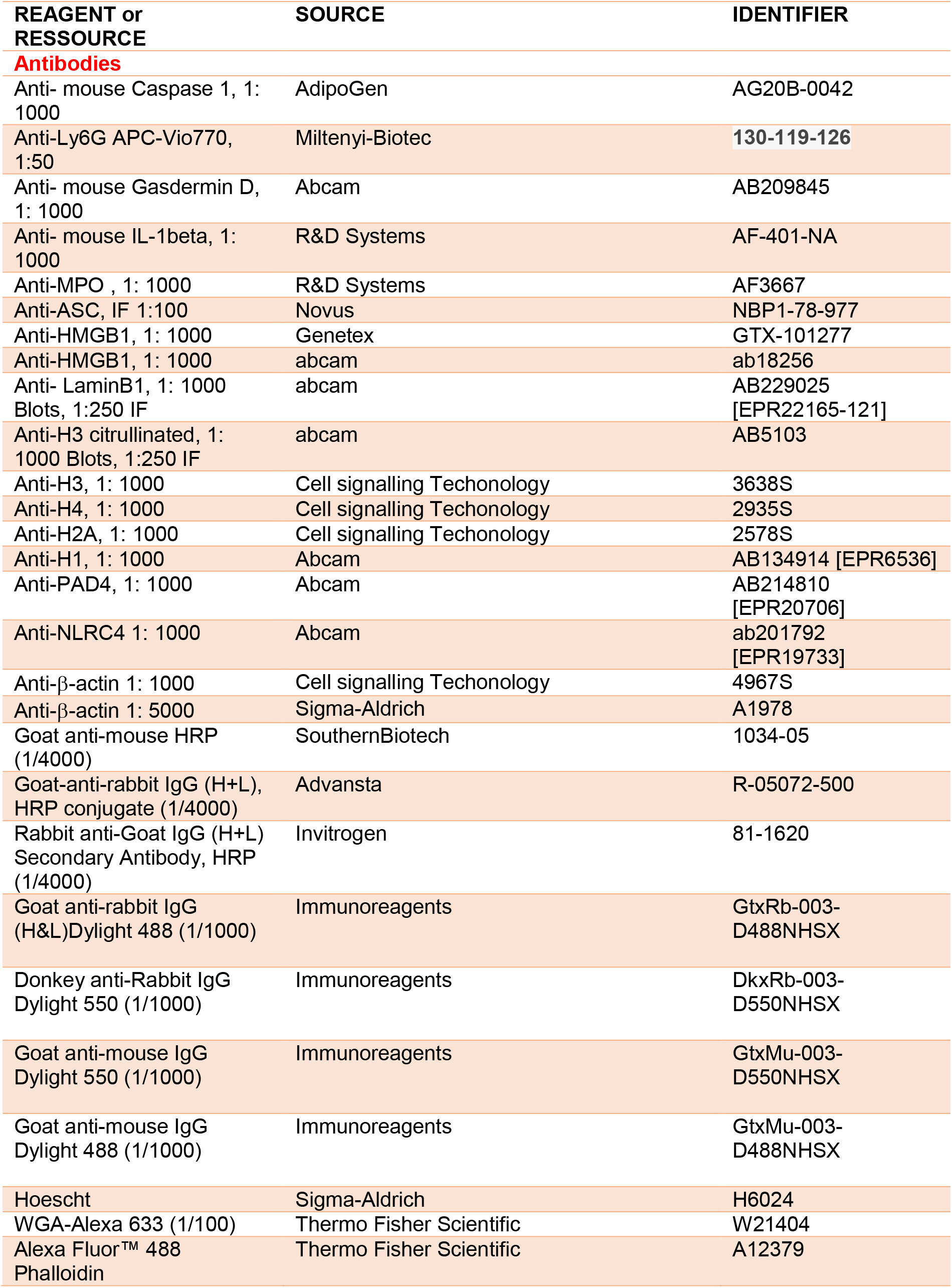
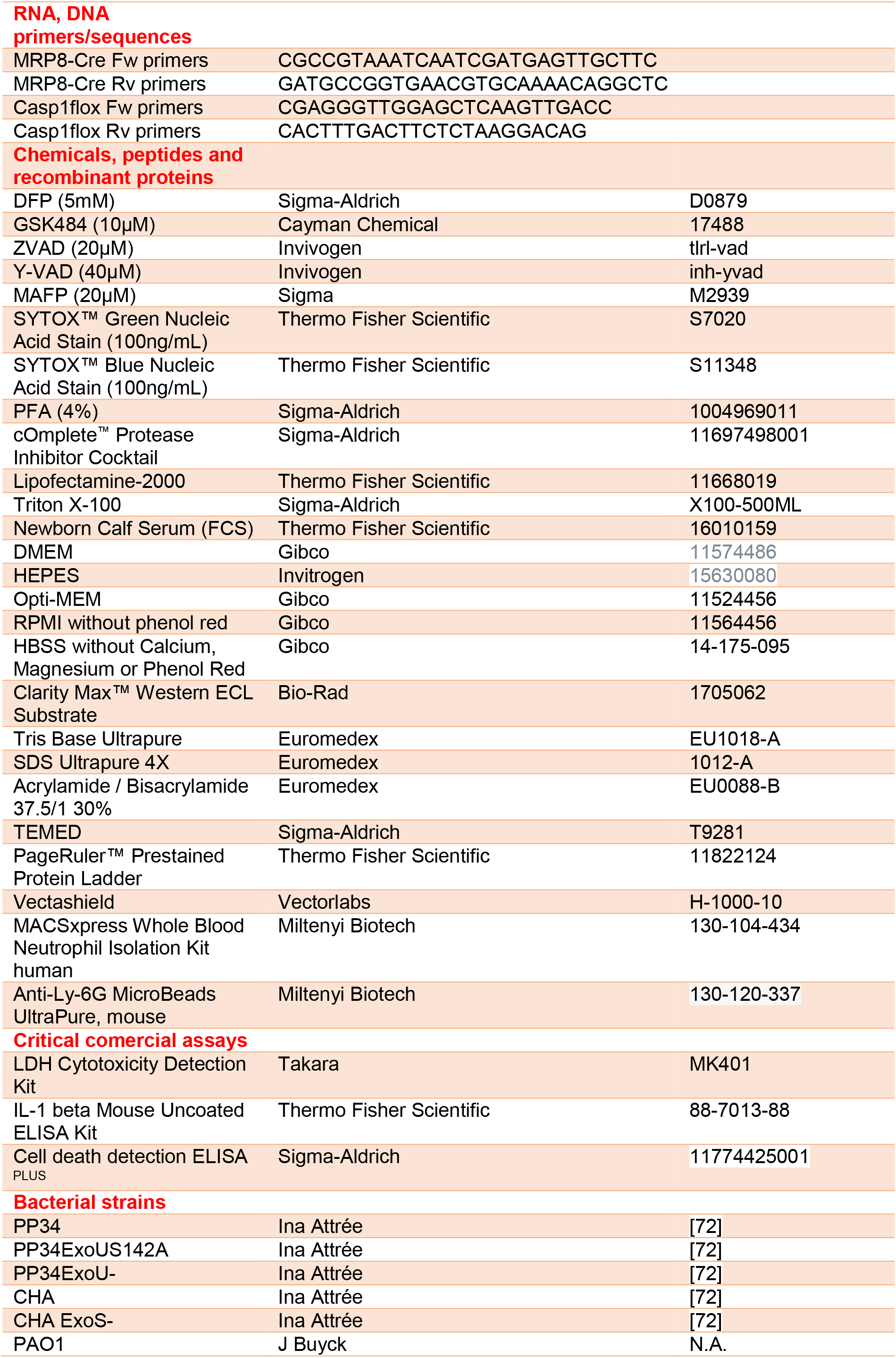
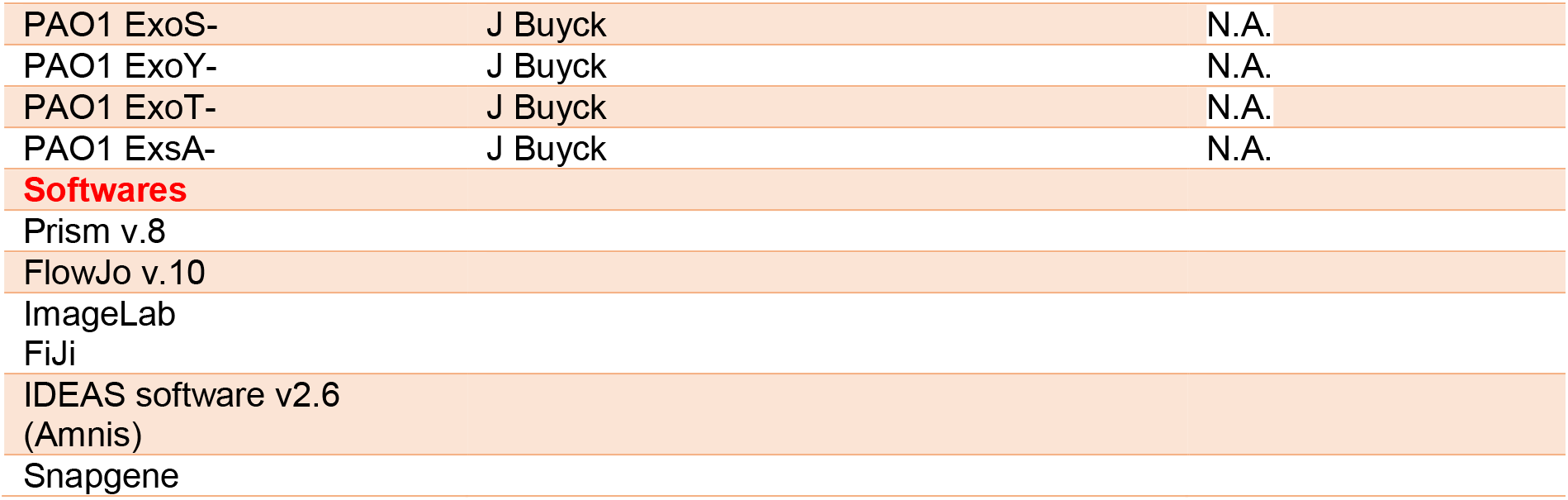
Reagents and Tools are available upon request to

